# Prebiotically Plausible Peptides can Self-assemble into β-rich Assemblies

**DOI:** 10.1101/2025.11.09.687475

**Authors:** Mikhail Makarov, Robin Kryštůfek, Matúš Friček, Edgar Manriquez-Sandoval, Soumya Dutta, Ján Michael Kormaník, Tadeáš Kalvoda, Václav Verner, Lucie Bednárová, Tatsiana Charnavets, Michal Lebl, Sean M. Brown, Romana Hadravová, Jan Konvalinka, Abhishek Singharoy, Stephen D. Fried, Klára Hlouchová

**Affiliations:** Department of Cell Biology, Faculty of Science, Charles University, Prague, 12800, Czech Republic; Institute of Organic Chemistry and Biochemistry, Czech Academy of Sciences, Prague, 16610, Czech Republic; Department of Physical Chemistry, Faculty of Science, Charles University, Prague 12843, Czech Republic; T. C. Jenkins Department of Biophysics, Johns Hopkins University, Baltimore, MD 21218, USA; School of Molecular Sciences, Arizona State University, Tempe, AZ, USA; Institute of Biotechnology of the Czech Academy of Sciences, BIOCEV, Vestec, 25250, Czech Republic; Department of Biological Sciences, University of Maryland Baltimore County, Baltimore, MD 21250, USA; Department of Chemistry, Johns Hopkins University, Baltimore, MD 21218, USA

**Keywords:** Peptide assembly, prebiotic chemistry, catalysis, protein structure, language models, peptide libraries, random sequences

## Abstract

Peptides can self-assemble into diverse morphologies in a programmable manner and hence are privileged building blocks used widely in nanotechnology. Most reported peptide nanostructures consist of one (or a few) defined sequence(s), as self-similarity is presumed to be essential for promoting assembly. While oligomerisation is seen as an important feature of the earliest functional polymers during the origin of life, prebiotic peptides were likely short, statistical, and non-templated – traits that seem incommensurate with robust self-assembly. Here we show that random 25-mer peptides can efficiently and spontaneously form highly thermostable, soluble assemblies rich with β-sheets. Notably, these nanostructures only emerge when random peptides are constructed with an “early” alphabet, consisting of the 10 canonical amino acids that were also prebiotically abundant – but not other alphabets tested. Hence, our findings show that peptide self-assembly does not require purity, and in fact compositional complexity is adaptive for preventing formation of insoluble structures. Altogether, this study showcases that unevolved sequences of prebiotically-abundant amino acids can readily produce foldable self-assembling polymers, thereby providing a potential steppingstone toward the first proteins, prior to the onset of purifying selection.

**Significance Statement:** Modern proteins are remarkable polymers that use a 20-letter amino acid alphabet arranged into specific sequences that arose from billions of years of evolution. However, in the prebiotic stages of Earth’s history, (i) some amino acids were likely not abundant – including all the canonical basic amino acids – as opposed to the canonical acidic amino acids, and (ii) sequences of peptides were likely statistical, preceding templated synthesis.

Here we report that random sequences composed from a prebiotically plausible acidic amino acid alphabet stand out in their capacity to promote peptide self-assembly. We observe that such soluble assemblies are β-sheet rich and may thus represent plausible scaffold of prebiotic interactions and catalysis.

## Introduction

Biochemistry researchers and students alike have long asked, Why did life since the last universal common ancestor (LUCA) ‘select’ the twenty α-amino acids as the established language of building proteins and peptides(1–5)? The groundbreaking work of Miller and Urey (6, 7) in the 1950s provided early evidence that unsupervised gas phase reactions under reducing conditions could have provided abundant access to some α-amino acids early in Earth’s history. Even at this incipient stage in the field of prebiotic chemistry, an apparent asymmetry was noted: the modern acidic amino acids – L-aspartic acid and L-glutamic acid – were generated in high yields, whereas the modern basic amino acids – L-arginine, L-histidine, and L-lysine – were not detected. The absence of these essential building blocks was rightly perceived as a roadblock for an early emergence of functional proteins (2), given that extant proteins are comprised of a balanced mix of acidic and basic residues, and basic residues are almost always involved in interactions with nucleotides and nucleic acids (2, 8). Analyses of the amino acids that are carried to Earth by carbonaceous meteorites (9) as well as recent examination of the Bennu and Ryugu asteroids (which were not subjected to terrestrial exposure) (10, 11) have only reinforced the view that the modern acidic amino acids were plentiful whereas the modern basic amino acids were unavailable in prebiotic sources. The overlap between the amino acid compositions from gas-phase atmospheric chemistry and extraterrestrial cosmic material have led to propositions of a consensus ‘early’ amino acid alphabet (12) that comprised of ten α-amino acids: alanine (Ala), aspartic acid (Asp), glutamic acid (Glu), glycine (Gly), leucine (Leu), isoleucine (Ile), proline (Pro), serine (Ser), threonine (Thr), and valine (Val) (Figure 1, grey and red circles).

**Figure 1.**
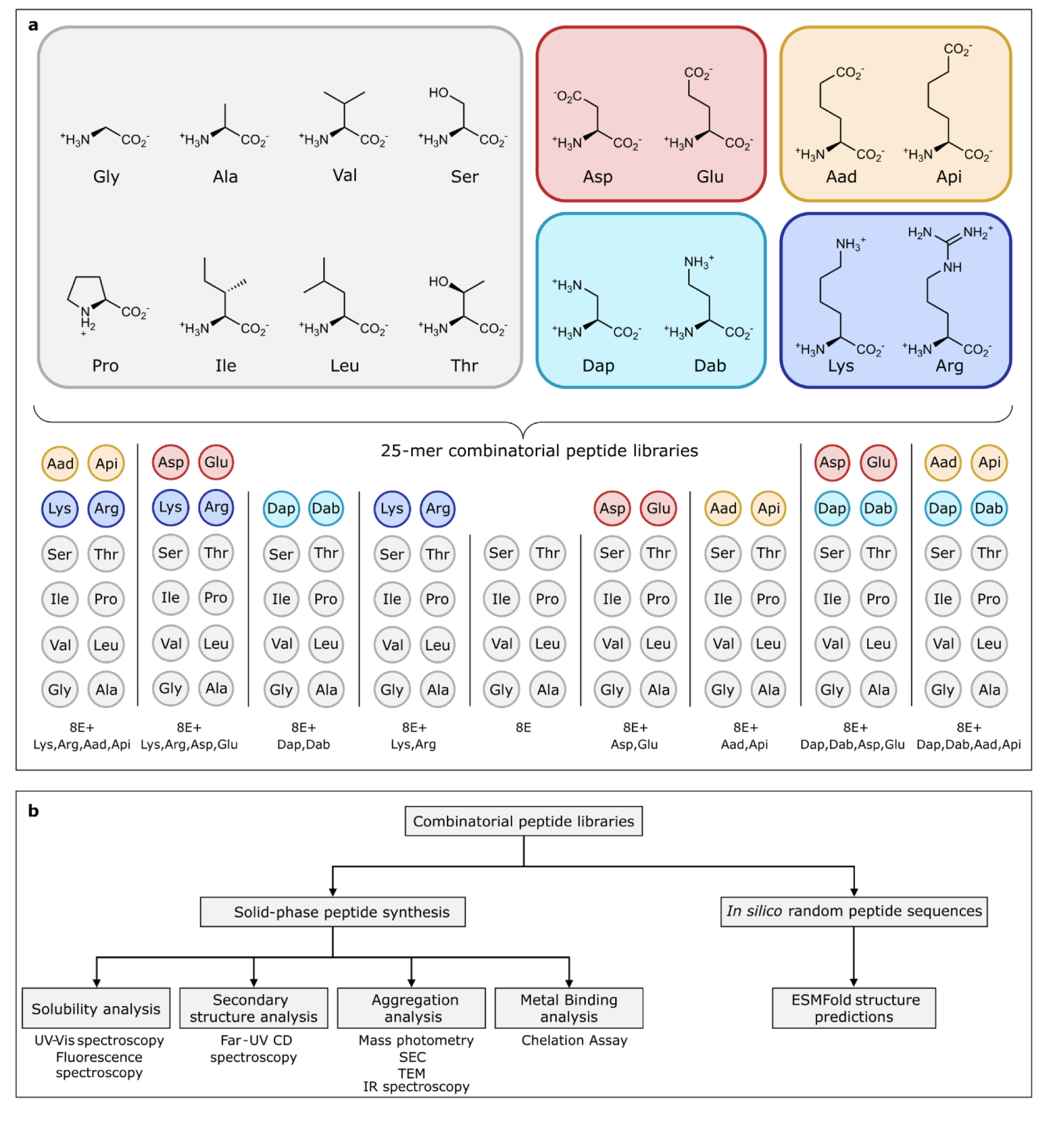
Design and evaluation of random peptide libraries using distinct amino acid alphabets. (**a**) Chemical structures of 16 α-amino acids assembled through solid phase peptide synthesis to generate random 25-mer peptides utilising one of nine distinct alphabets (collections of amino acids). Dap, 2,3-diaminopropionic acid; Dab, 2,4-diaminobutyric acid; Aad, 2-aminoadipic acid; Api, 2-aminopimelic acid. (**b**) Methods used in this study to interrogate random peptide libraries.

Whether such a reduced chemical repertoire could have served as a practical inventory for building up the earliest biopolymers is not apparent, given the essential roles aromatic, basic, and sulfur-containing amino acids play in modern protein biochemistry. Recent efforts from us and others (13, 14) have at least substantiated this possibility by showing that these ‘early acidic’ proteins can adopt conformations with secondary structure, can form strong associations with RNA mediated by divalent cation coordination (15), and strikingly, are much less dependent on chaperones to adopt soluble stable conformations (14, 16, 17). A recent report has shown that distinct small protein folds can be reliably designed with the reduced alphabet using the modern de novo design methods (18). On the other hand, the absence of positively charged residues in the early alphabet implies these peptides cannot form salt-bridges. This would seem to make the early acidic proteins deficient in their capacity to self-assemble and raises the possibility that alternative alphabets that include prebiotically available basic residues like L-2,3-diaminopropionic acid (Dap) and L-2,4-diaminobutyric acid (Dab) (Figure 1, cyan circles) could possess some adaptive advantages (19).

Here, we extensively characterise nine random peptide libraries (heterogenous mixtures of sequences), each composed of various combinations of acidic and basic amino acids (some canonical, some prebiotic, and some counterfactual, Figure 1), to interrogate effects on solubility, folding propensity, and self-assembly potential. The data shows that acidic alphabets are remarkable in their capacity to promote folding as well as oligomer assembly mediated through β-sheets. We find unambiguously that any inclusion of basic amino acid residues – either in isolation or alongside acidic residues – diminishes these properties. The assemblies formed by the prebiotically plausible acidic peptides are quite thermostable but do not resemble amyloid structure (based on both solubility and morphology). Altogether, these findings point to acidic polypeptides as a potential intermediate in the early evolution of peptides and proteins. Though alien in some ways compared to today’s charge-balanced polypeptides, we observe that the early acidic alphabet could provide key advantages that facilitated the initial emergence of biopolymers.

## Results

### Synthesis of Charge-Variant Libraries

Based on the presumption of prebiotic feasibility of non-templated peptide synthesis (21–24), libraries of 25-mer peptides, constructed as random combinations of amino acids within the library, were synthesized. This was performed by standard Fmoc-based solid phase peptide synthesis using the isokinetic mixtures as previously reported (20) (see Methods and Supplemental Data S1). Each library sampled one of nine different possible alphabets (Figure 1a), which could comprise either 8, 10, or 12 constituent α-amino acids. The common core of all libraries (library “8E”) only uses the eight uncharged canonical and prebiotic amino acids (grey), which we synthesised as a control. We considered two basic alphabets: 8E+Lys,Arg adds the two modern basic amino acids with long side chains (dark blue), whereas 8E+Dap,Dab adds the two most prebiotically available basic amino acids (cyan). We considered two acidic alphabets: 8E+Asp,Glu which adds the two canonical and prebiotically plausible acidic amino acids (red), and 8E+Aad,Api (L-α-2-aminoadipic acid, L-α-2-aminopimelic acid; orange) which is a ‘counterfactual’ alphabet. These amino acids were not prebiotically abundant but mimic the longer side chains of Lys and Arg and provide orthogonal means to address whether Asp and Glu provide unique advantages over hypothetical alternatives. Finally, we also considered four mixed alphabets in which acidic and basic amino acids coexist with varying side chain lengths.

We established the quality of these libraries with both MALDI mass spectrometry (Supplemental Figure S1) and HPLC amino acid analysis (Supplemental Figure S2, measurements compiled in Supplemental Data S2). MALDI showed distributions of precursor masses quite similar to theoretical predictions for each library (Supplemental Figure S1). For instance, the predicted average molecular weight for a 25-mer peptide in the 8E+Dap,Dab library is 2330 Da; we observe that it is 2288 Da (difference 42 Da or ∼0.44 amino acids), which is similar for the other libraries. Amino acid analysis showed correct incorporation of the amino acids in each library and close-to-uniform distributions in the amino acids’ stoichiometry (Supplemental Figure S2). For instance, since the 8E+Lys,Arg,Aad,Api library has 12 amino acids, each amino acid should represent ∼8.3% of the total amino acid content theoretically; we observe that each amino acid is present at 8 ± 3% of the content, and this is similar for the other eight libraries. We conclude that the isokinetic mix strategy can produce reasonably representative random peptide libraries, even spanning those with distinct isoelectric points and noncanonical amino acids.

An array of biophysical and computational methods was then used to characterize the libraries (Figure 1b).

### Solubility Profiles of Charge-Variant Libraries

Lyophilised peptide libraries were resuspended in aqueous Britton-Robinson (ABP) buffers of various pH’s and supplemented with NaCl to either a medium (50 mM) or high (500 mM) ionic strength. After mixing, the solutions were centrifuged at 21,000 x g for 30 min to remove insoluble material, and the solubility of the alphabets were assessed by measuring peptide concentration in the soluble fraction by UV-Vis absorption spectroscopy (Supplemental Figure S3, measurements compiled in Supplemental Data S2).

Similarly to what we previously observed in 8E+Asp,Glu (20), the acidic 8E+Aad,Api alphabet showed poor solubility at low pH, and high solubility at neutral and high pH (Supplemental Figure S3). These results can be rationalised since carboxylic acids are expected to be protonated and neutral at pH below 5, thus promoting aggregation. To our surprise, all other six libraries that contained basic amino acids were highly soluble at all pH’s (Supplemental Figure S3), when we might have expected 8E+Lys,Arg and 8E+Dap,Dab to mirror the acidic libraries and be insoluble at higher pH’s (although the pK_A_ of Arg is further away from physiological pH).

In general, ionic strength has minimal effect on solubility, with the only exception being the acidic alphabets (8E+Asp,Glu and 8E+Aad,Api) at physiological pH, where high salt concentration is deleterious (Supplemental Figure S3). In other words, it is only the acidic alphabets that evidence the Hofmeister salting-out effect typically ascribed to globular proteins (25, 26). This result suggests that the acidic alphabets have a rather unique balance of hydrophobicity and hydrophilicity, which can promote aggregation under some conditions (i.e., low pH, high ionic strength) and potentially folding under other conditions (i.e., neutral pH, medium ionic strength).

### Soluble Aggregate Formation in Charge-Variant Libraries

Size exclusion chromatography (SEC) can provide an orthogonal means to assess aggregation, and unlike centrifugal pelleting assays, which measure precipitating aggregation, SEC is sensitive to the formation of soluble oligomers/aggregates. We examined all the peptide libraries in the pH range between 3–11, finding that in all cases, the majority of the peptide mass eluted as a single peak with an elution volume between 13–16 ml (Figure 2). These data indicate that most of the soluble peptides are monomeric and do not form oligomers or soluble aggregates. The two acidic libraries (8E+Asp,Glu and 8E+Aad,Api) showed the greatest pH-dependence on their elution profile in a predictable fashion. At pH 11, the peptides eluted at 13 ml, implying larger hydrodynamic radii when the molecules are the most polyanionic (and most likely unfolded). At pH 3, where these peptides are more neutralised and higher levels of precipitation are observed (cf. Supplemental Figure S3), the soluble peptides eluted at 16 ml, implying more compact structures with lower hydrodynamic radii. Curiously, the two basic libraries (8E+Lys,Arg and 8E+Dap,Dab) did not show the expected reciprocal behaviour of becoming more compact at high pH, where the peptides would become more charge neutralised. Instead, the position of the primary peak remained at ca. 16 ml, though the chromatographic feature does broaden and narrow in a pH-dependent manner (Figure 2). The SEC profiles of the four libraries with both acidic and basic residues responded to pH in a similar, if slightly attenuated, way as the acidic libraries.

**Figure 2.**
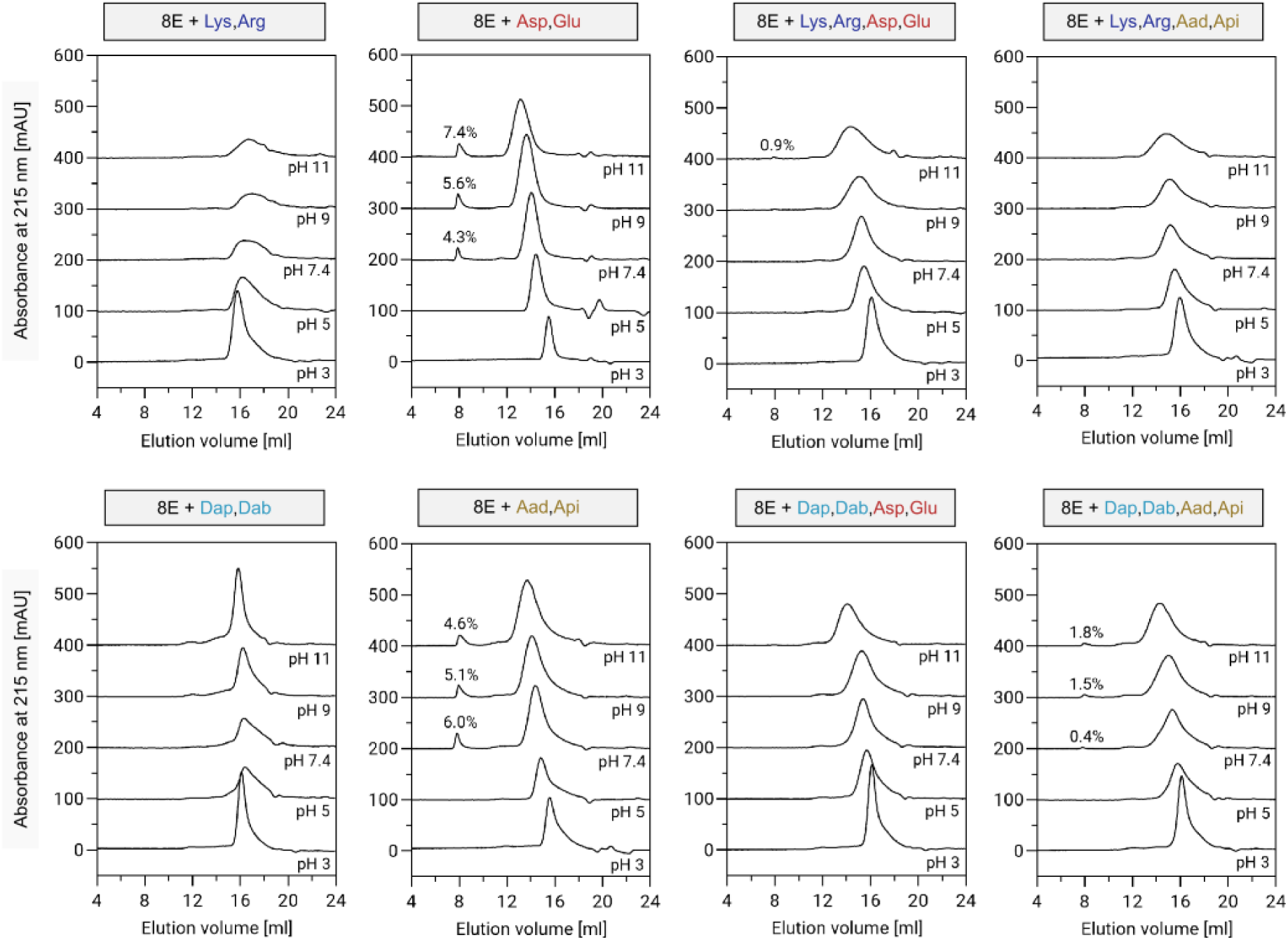
The aggregation propensity of eight peptide libraries. Aggregation propensity of the indicated libraries was measured in 20 mM ABP buffer at different pH’s and at medium ionic strength (50 mM NaCl) with size-exclusion chromatography (SEC). SEC chromatograms report absorbance at 215 nm as a function of elution volume. The data for 8E+Asp,Glu library were taken from previous work (20).

We detected soluble aggregates forming as a separate feature at low elution volume (8 ml), but *only* in the two acidic libraries and *only* under pH’s where these libraries did *not* produce precipitating aggregates (Figure 2 and Supplemental Figure S3). This is a key observation as self-assembly into larger structures has been posited as an important property for prebiotic peptides (27, 28).

These experiments were repeated at high ionic strength (500 mM NaCl, Supplemental Figure S4), which produced SEC chromatograms that were largely similar but differed in two key respects. Firstly, the pH-associated changes to the peak position were reduced, which is reasonable because the salt screens repulsive ion-ion interactions that would promote peptide chain radius expansion. Secondly, and more importantly, the soluble aggregate feature we detected in 8E+Asp,Glu is greatly diminished at 500 mM NaCl, implying that these assemblies involve electrostatic interactions that are screened out at high ionic strength. Notably, the ability of 8E+Asp,Glu peptides to form assemblies is strongly dependent on their length, with assembly formation not observed for 8- and 15-mer versions of the library (Supplemental Figure S5). This indicates that a critical peptide length is necessary for random peptides to form higher-order architectures.

### Beta Sheets Mediate Soluble Assembly of Prebiotic Acidic Peptides

To interrogate the nature of the peptide assemblies formed in the 8E+Asp,Glu library, we performed SEC preparatively and isolated the early-eluting peaks (Figure 3). The peptide mixture was analysed by CD spectroscopy, and remarkably, this subpopulation evinced clear signature for β-sheets, including a clear negative peak at 215 nm, and positive peak at 192 nm. When the subpopulation was hydrolysed and analysed with amino acid analysis, we saw an enrichment for the aliphatic branched amino acids, such as Val, Leu, and Ile, and depletion of Pro and the charged amino acids, Asp and Glu. This apparent sensitivity of the assembly formation to the library composition was further confirmed by synthesizing the 8E+Asp,Glu library with individually skewed amino acid ratios (Supplemental Figure S6 and Table S1). While increasing the level of Ile was particularly beneficial for enhancing formation of assemblies, increasing the levels of Pro, Glu, or Asp abrogated assembly formation.

**Figure 3.**
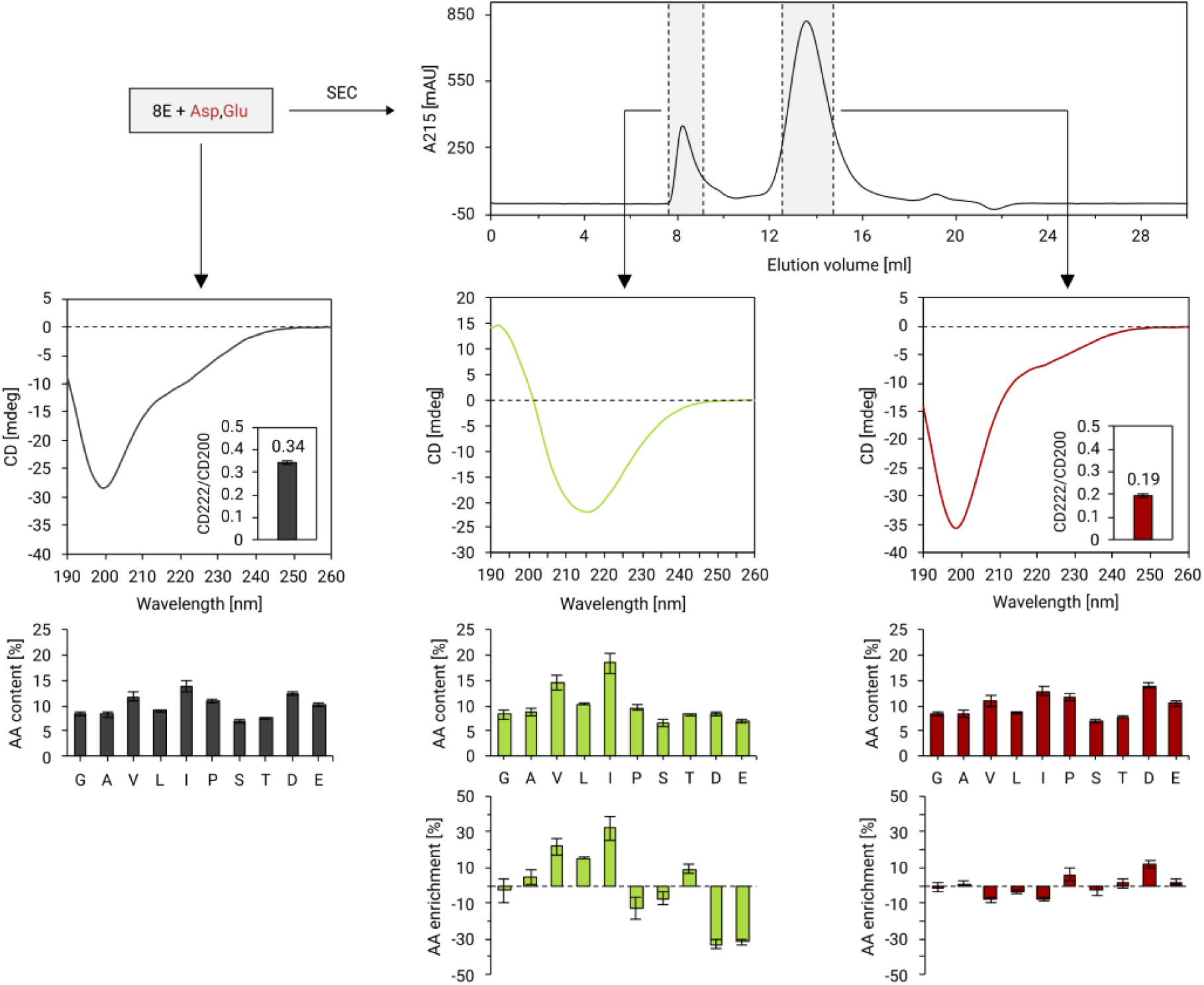
The resolution of a β-sheet rich subpopulation of peptides from the soluble aggregate fraction of the 8E+Asp,Glu library. The 8E+Asp,Glu library was dissolved in 10 mM ABP buffer (pH 7.4) and resolved by size-exclusion chromatography. The soluble aggregate fraction (lime) at 8.5 ml was isolated and analyzed by far-UV circular dichroism (CD) and HPLC amino acid analysis. The far-UV CD spectrum possesses a negative peak at 215 nm and a positive peak at 192 nm, signaling β-sheet secondary structure. The HPLC amino acid analysis showed enrichment in branched amino acids (Val (V), Leu (L), and Ile (I)) and depletion of Pro (P) and acidic residues (Asp (D), Glu (E)). The monomer fraction (red) at 13.5 ml possesses a far-UV CD spectrum and amino acid content similar to that of the input peptide library (blue).

This dependence of assembly formation on amino acid frequencies matches closely with the amino acids known to have higher (Val, Leu, Ile) and lower (Glu) propensity to occur in β-sheets (29). On the other hand, the late-eluting peak, which we can assign to monomeric peptides, possessed a CD spectrum and an amino acid composition relatively similar to the original unseparated peptide library (Figure 3), though with an even more negative peak at 200 nm (corresponding to loops and coils) and a narrower shoulder at 222 nm (corresponding to regular secondary structures, α-helices primarily).

The remarkable overall folding potential of the 8E+Asp,Glu library comes into starker focus when compared with the other libraries (Supplemental Figure S7, spectral data compiled in Supplemental Data S2). Even the chemically similar 8E+Aad,Api alphabet shows a diminished spectral feature at 215 and 222 nm suggesting peptides with attenuated secondary structure when the negatively charged carboxylic acids are placed further away from the backbone. Moreover, we find that all six alphabets containing basic residues, either alone or in combination with acidic residues, exhibit strikingly diminished capacity to form secondary structures (Supplemental Figure S7). We were surprised by these results particularly in relation to the four mixed-charge libraries, as they have isoelectric point distributions more representative of naturally evolved proteins. These findings are invariant to pH, and indeed it is only the acidic alphabets whose secondary structure capacity show any pronounced dependence on pH. As a control to ensure that the folding properties were not influenced by the ABP buffer (which enabled the interrogation of a broad pH range), we recorded CD spectra for 8E+Asp,Glu, 8E+Lys,Arg, and 8E+Lys,Arg,Asp,Glu in Tris-HCl (pH 7.4) and found this change resulted in no significant differences with respect to ABP buffer (Supplemental Figure S8). As a further control, we acquired CD spectra of these random peptide libraries with varying amounts of 2,2,2-trifluoroethanol (TFE) added (between 10-90% (v/v)) as a cosolvent in order to induce α-helical structure (Supplemental Figure S9). All the alphabets responded to TFE induction, with pronounced negative features at 205 and 222 nm appearing in all cases with TFE concentrations of 50% and higher. These experiments serve as useful controls that show all libraries have, in principle, the capacity to fold, and none of the noncanonical amino acids impose steric constraints that explicitly prevent it.

### The peptide assembly is thermally stable but distinct from amyloid structures

We used mass photometry to estimate the aggregation number of the peptide assemblies in the 8E+Asp,Glu library. These experiments identified a non-Gaussian peak with a maximum at ca. 80 kDa (ca. 32 peptides, Supplemental Figure S10b) and a fat tail extending to 400 kDa, which is absent entirely in the late-eluting SEC fraction (Supplemental Figure S10c) and enriched substantially in the early-eluting SEC fraction (Figure 4a). Hence, we conclude that the 8E+Asp,Glu alphabet admits a rather intriguing characteristic: a sub-population of peptides that spontaneously assemble into larger soluble structures consisting primarily of β-sheets. Most intriguingly, this property emerges in random sequence space, though only with the appropriate alphabet.

**Figure 4.**
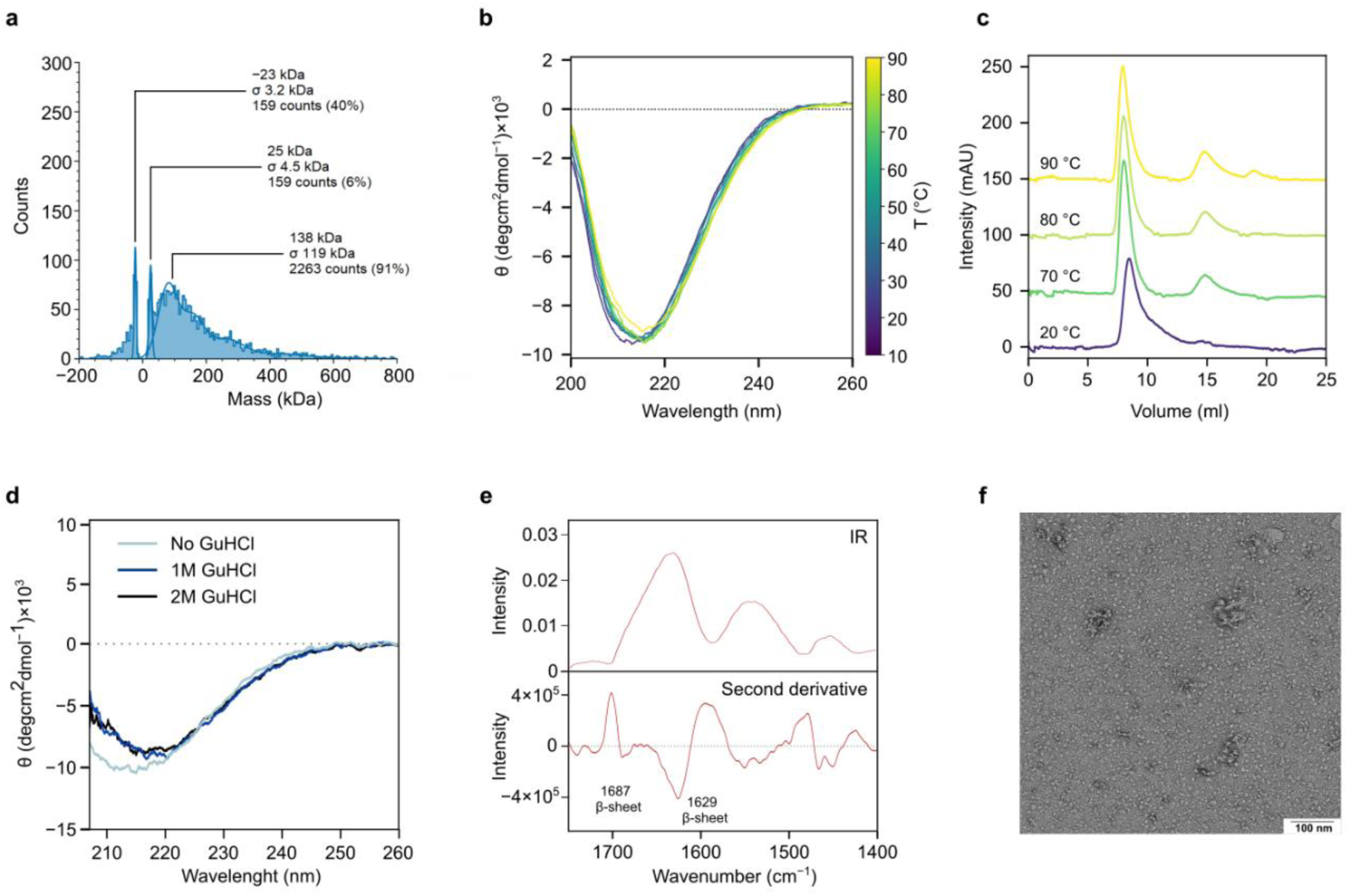
Size, morphology and stability of the 8E+Asp,Glu assemblies. (**a**) The mass histogram of the isolated assembly fraction of 8E+Asp,Glu library in 10 mM Tris-HCl (pH 7.4), 50 mM NaCl assessed by mass photometry. (**b**) Temperature dependent far-UV circular dichroism (CD) of assembly fraction in 10 mM ABP (pH 7.4) performed in temperature range shown on scale in panel c. (**c**) Size exclusion chromatography of isolated assembly fraction performed in 10mM ABP (pH 7.4) after 1 hour incubation at 20, 70, 80 and 90 °C. ( **d**) Far-UV CD of assembly fraction in 10mM ABP (pH 7.4), with increasing concentration of guanidine hydrochloride (Gu*HCl) as indicated in the panel. (**e**) Infrared spectrum of the 8E+Asp,Glu library and spectrum derivative with assigned spectral bands of interest. (**f**) Transmission electron micrograph of the isolated assembly fraction of 8E+Asp,Glu library. The electron micrographs were obtained on a sample with 20 μg/ml concentration in 10 mM Tris-HCl (pH 7.4).

To evaluate the thermal stability of the 8E+Asp,Glu assemblies, we analyzed the isolated assembly fraction using temperature-dependent CD spectroscopy. The analysis revealed that the characteristic β-sheet structure of the purified assemblies was retained up to 90 °C, showing remarkable resistance to thermal denaturation (Figure 4b). This observation was further supported by size exclusion chromatography with the purified assemblies performed right after 1 hour incubation at 70 °C, 80 °C, and 90 °C, which showed only minor signs of disassembly (Figure 4c). Similarly, the assemblies are resistant to denaturation at modest concentrations of GuHCl (Figure 4d). The infrared spectrum of 8E+Asp,Glu library (Figure 4e) also admits signatures associated with assembly: an amide I band centered at 1629 cm^−1^ in combination with a shoulder at 1687 cm^−1^, indicating substantial β-sheet population.

Additionally, we collected transmission electron micrographs of the 8E+Asp,Glu assemblies and compared them with the monomer SEC fraction and total fractions of the 8E+Asp,Glu and 8E+Asp,Glu,Lys,Arg libraries (Figure 4f). Globular particles were observed in the assembled peptide species (detectable in the total 8E+Asp,Glu fraction and enriched in the early-eluting SEC fraction), but no fibrillar structures that would potentially resemble amyloids were observed in any micrographs collected.

### Charge-variant libraries bind metal ions with different preferences

Recent theoretical and experimental studies have suggested that metal ions may have played an important role during the earliest stages of protein evolution by stabilizing primitive acidic peptides and expanding their structural and functional repertoire before the emergence of modern protein folds (28). Such a scenario is particularly attractive for acidic peptide alphabets, which lack the positively charged residues characteristic of modern proteins yet possess abundant carboxylate groups capable of coordinating metal ions. Because we found that random peptides constructed from the prebiotically plausible acidic alphabet spontaneously assemble into highly stable β-sheet-rich oligomers, we next asked whether these assemblies also display distinctive metal-binding properties. To address this question, we measured binding of biologically relevant divalent metal ions (Fe²⁺, Ca²⁺, Zn²⁺, Ni²⁺, Co²⁺) using the metal-sensitive dye Fura-2 in the presence and absence of selected peptide libraries (Figure 5). Libraries 8E+Asp,Glu and 8E+Asp,Glu,Lys,Arg were used to evaluate the influence of differently charged residues on metal binding, whereas the 19F library - representing the complete set of amino acids (excepting Cys, excluded for synthetic reasons) - served as a control that included additional potential ligands such as histidine.

**Figure 5.**
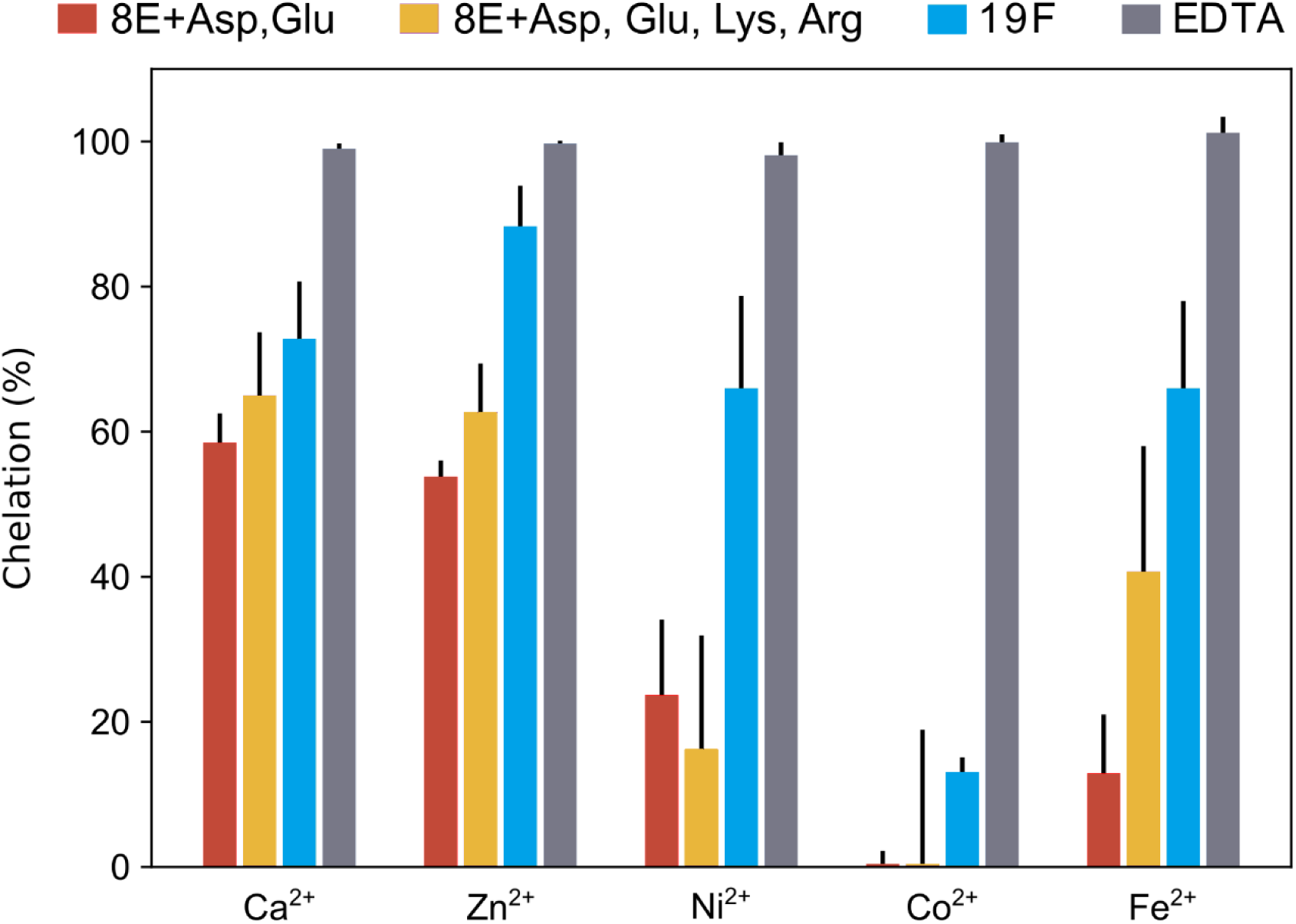
Metal ion binding propensity. Chelation of biologically relevant metal ions (Ca^2+^, Zn^2+^, Ni^2+^, Co^2+^, and Fe^2+^) was measured for three canonical amino acid alphabets (where 19F stands for the full set of canonical amino acids excepting Cys) using a fluorescent reporter-based assay. EDTA was used as a positive control. Error bars indicate the standard deviation from triplicate measurements.

All random-sequence libraries exhibited measurable metal-binding propensities, which increased with the compositional complexity of the libraries (Figure 5). Among the tested ions, 8E+Asp,Glu displayed the strongest affinity for Ca²⁺, consistent with Hard and Soft Acids and Bases (HSAB) theory (30), and this binding was enhanced by the addition of further amino acid types. Notably, Asp and Glu are also the most frequent Ca²⁺-binding residues in modern proteins (31). In contrast, Zn²⁺ - an ion intermediate between hard and soft acids – showed unexpectedly strong binding to the 8E+Asp,Glu library, with further enhancement observed for the 19F library, likely reflecting the contribution of histidine. The substantial Zn²⁺ affinity of 8E+Asp,Glu is intriguing given that stable coordination of transition metals in modern proteins typically requires softer donor groups (31). As anticipated from HSAB principles, binding of Fe²⁺, Ni²⁺, and Co²⁺ was limited for 8E+Asp,Glu but markedly increased in the 19F library.

### Large Language Models Recapitulate Structure-Forming Potential of Acidic Residues

Obtaining structural information on our random peptide libraries is challenging on two levels: (i) the vast compositional diversity of the libraries (essentially every sequence in the mixture is unique) and (ii) these short unevolved 25-mer peptides are expected to be dynamic and conformationally heterogenous. Recently, large language models have been developed for protein structure prediction (32, 33). These models do not rely on multiple sequence alignments, and hence it has been argued that they have learned a more generic relationship between sequence and structure that is not dependent on biological sequences or even biological amino acid compositions. We therefore carried out ESMFold structure predictions on 50,000 unique and randomly generated 25-mer sequences from four alphabets comprising only canonical amino acids (8E, 8E+Asp,Glu, 8E+Lys,Arg, and 8E+Lys,Arg,Asp,Glu), as ESMFold does not support noncanonical amino acids.

In consonance with our experimental studies, ESMFold predicts that peptide compositions drawing from the prebiotically plausible 8E+Asp,Glu alphabet are slightly more compact (median *R_g_* = 1.63 nm) than peptides drawing from the basic alphabet (median *R_g_* = 1.82 nm) or the charge-balanced alphabet (median *R_g_* = 1.69 nm) (Supplemental Figure S11a). On the other hand, ESMFold predicts that the 8E library, which possesses no charged residues, generates the most compact structures (Supplemental Figure S11a, median *R_g_* = 1.36 nm). At face value, this result appears to be in direct contrast with CD spectroscopy (Supplemental Figure S11b) of this peptide library, which evidences very little secondary structure. However, fluorometric solubility assays show that the 8E library has poor solubility at all pH’s (Supplemental Figure S11b), which can be ascribed to its very hydrophobic composition (results for all other assays on the 8E peptide library are given in Supplemental Figure S12). One can reconcile these findings by suggesting that 8E *does* possess many sequences that can fold in principle, but those peptides also aggregate and are therefore depleted from the soluble fraction on which CD spectra were acquired.

## Discussion

Our research into the properties of prebiotically-plausible amino acid alphabets began with the assumption that a severe reduction in the chemical inventory would imply proteins with more limited folding and function compared to modern proteins with their access to aromatic, basic, amidic, and sulfur-containing building blocks. Although such a rudimentary alphabet has been shown to meet simple requirements – such as the ability to fold (18, 20), bind RNA (15), or bind organic cofactors (40) – its potential role before the advent of templated synthesis has remained unclear. In light of the results presented here, the early 8E+Asp,Glu alphabet appears to constitute a uniquely adaptive pool of building blocks, well suited to act as a transitional pool toward early biopolymers. These adaptive features stem from the ability of statistical peptides of this composition to self-assemble and coordinate metal ions.

Among the tested charge-variant libraries, 8E+Asp,Glu possesses substantial secondary structure propensity whilst also maintaining high solubility, a delicate “balancing act” that it pulls off, unlike the hydrophobic 8E alphabet and the modern alphabet (cf. Supplemental Figures S11-12 and previous work (14, 20)), which are highly prone to aggregate. The fact that the 8E+Aad,Api alphabet recapitulates most of the key biophysical features of 8E+Asp,Glu shows that there is *some* leeway in the chemical space that admits peptides that can fold and assemble. These results suggest that side-chain length is perhaps less critical of a modulator than we had initially hypothesised (20). Together, these findings suggest that 8E+Asp,Glu was a strong candidate for assisting the emergence of proteins prior to purifying selection. The modern 20-letter alphabet has great functional breadth, but this is realized through specific sequences that have been finessed by selection and frequently rely on chaperones to fold (14, 41). Random sequences utilising this more advanced amino acid repertoire, on the other hand, suffer from aggregation. An amino acid set that promotes high solubility and foldability across unevolved sequences would have played a key adaptive role during the early stages of life’s emergence, for which the early 8E+Asp,Glu alphabet would be more suitable.

Both experiments and AI modelling agree that β-sheets are rare amongst random peptide sequences as monomers. We report here, however, that a subset of 8E+Asp,Glu peptides self-assemble into higher-order oligomers with pronounced β-sheet content. This finding is particularly germane to the theory that amyloids could have played a key role during the origin of life (the amyloid world hypothesis), with their ability to form complex structures and propagate using short peptides as monomers (27). Nevertheless, the formation of amyloids typically depends on specific, repetitive sequence patterns—features that would have been difficult to achieve under prebiotic conditions.

Compositionally amphiphilic peptides (similar to those found in the 8E+Asp,Glu library) have been explored by design in the field of peptide nanotechnology, assembling into supramolecular structures rich in β-sheets (42). Our results here constitute, to the best of our knowledge, the first observation of soluble oligomeric β-sheet assemblies arising from *random* and *highly heterogenous* peptide sequences comprising early amino acids, i.e. sequences that would plausibly be formed preceding templated ribosomal synthesis and the onset of cellular life. Whilst we would refrain from referring to these species as fibrils or amyloids (their molecular weights span between 80–400 kDa, they are soluble, and do not show fibrillar morphology in electron micrographs), they provide evidence that beta structures were also available to early peptides via oligomerisation. Most likely, soluble nanostructures form because the sample’s heterogeneity precludes a high enough number of mutually compatible peptides to be available to form larger insoluble structures. Importantly, self-assembled peptides stemming from relatively short sequences have been discussed as plausible sites of (prebiotic) catalysis and interaction scaffolds, in the absence of longer polypeptides and tertiary folding (42–47). Intermolecular assemblies of peptides lead to net release of water molecules at nonpolar interfaces and can lead to increased reaction rates (48). Besides the involvement of the non-polar amino acids (Val, Leu, and Ile, all of which are enriched in the assemblies) in stabilization of the peptide nanostructures, the intermolecular interaction between the β-sheets are likely further supported by the metal ions that bind substantially to the 8E+Asp,Glu library (such as Ca^2+^) as implied by the assembly weakening in high ionic strength. The self-assembling capacity of random 8E+Asp,Glu peptides emerges with peptides longer than 20 amino acids (Supplemental Figure S5). Such a length threshold could have represented a critical step in the transition from short oligomers towards structurally more defined, self-organizing peptide systems capable of exhibiting primitive functional properties. Moreover, the high thermal stability and stability to denaturant of the observed assemblies suggests that such β-sheet-based architectures could have endured the fluctuating conditions of the prebiotic environment, and provided them with a fitness advantage.

The lower affinity of Asp and Glu residues for transition metals, as predicted by the HSAB principle, aligns well with the reducing and alkaline nature of prebiotic environments, where ions such as Ni²⁺ or Fe²⁺ would have been scarce due to their tendency to precipitate under such conditions. Conversely, the binding of alkaline earth metals such as Ca^2+^ and Mg^2+^, which are both hard ions, may have promoted primitive catalytic activity and contributed to the emergence of the earliest metalloproteins, which were likely smaller and relied on simpler coordination environments compared to modern metalloproteins, which use Cys and His residues in evolutionarily optimized geometries.

A view of the protein universe based solely on extant proteins might suggest that polypeptide sequences which fold and assemble are rare, and once discovered, are conserved, thereby explaining the relatively small number of unique fold topologies used in Nature (49). Nevertheless, this is a view based on the modern amino acid alphabet. An early acidic amino acid alphabet can produce random polypeptides with these properties with higher probability. With this view in mind, the emergence of polymers that can fold and self-assemble from a primordial soup rich in α-amino acids begins to appear less of a statistical anomaly, but rather, quite plausible.

## Materials and Methods

### Solid-phase synthesis

Synthesis was conducted using the standard Fmoc peptide synthesis methodology using the SPENSER automated synthesizer (IOCB Prague, Czech Republic). The synthesis of the libraries was conducted on 384-well filter plates (Cytiva 5072-N) with a reaction scale of 1280 nmol per well. Incubations were carried out using an apparatus that alternated between N _2_ overpressure (0.05 atm) and suction (−0.75 atm) to mix the well contents, with a period of 5 s between 200 ms suction pulses. To start the synthesis, Fmoc-protected amino acids were weighed (see Supplemental Data S1), combined, and then dissolved to form a 0.3 M solution with 0.375 M OxymaPure in dimethylformamide (DMF) to form an isokinetic mixture. Deprotection was performed by sequential addition of 6 × 40 μL of 20% piperidine in DMF, with each addition incubated for 30 min. Wells were washed using 35, 80, and 6 × 35 μL of DMF. Supernatants were removed via suction applied for 1 min. Coupling was carried out using 2 × 45 μL of isokinetic mixture, and 375 mM N,N’-diisopropylcarbodiimide in DMF and incubated for 3 h each time, followed by washing, deprotection and washing. After the final deprotection cycle, the resin was thoroughly rinsed with DMF and all liquid contents were removed via suction applied for 1 min. After that, resin was taken out of the 384-well filter plate via centrifugation at 4000 × *g* for 5 minutes and combined. The dried, combined resin underwent treatment with 10 ml of Mixture K (trifluoroacetic acid–water–triisopropylsilane, 95:2.5:2.5, v/v) per gram of resin for 2 h. The resin was then filtered out, and the resulting peptide solution was precipitated with diethyl ether (20 ml/1 ml Mixture K) and centrifuged at 4000 × *g* for 5 minutes. The resulting pellet was suspended in 1 mM HCl and subjected to lyophilization. Before conducting subsequent experiments, all peptide libraries were further lyophilized thrice from 1 mM HCl overnight, establishing Cl^-^ as the counterion.

### Quality control

The molecular weight distributions of combinatorial peptide libraries were confirmed by mass spectrometry using UltrafleXtreme MALDI-TOF/TOF mass spectrometer (Bruker Daltonics, Germany) according to the standard procedure. Prior to the experiment, peptide libraries were dissolved in acetonitrile:water (1:1) mixture to the final concentration of 10 mg/ml. 2,5-Dihydroxybenzoic acid (DHB) was used as a matrix.

Prior to the amino acid analysis, the library samples were hydrolyzed in 6 M hydrochloric acid at 110 °C for 20 h and the hydrolysate was evaporated and reconstituted with 0.1 M hydrochloric acid containing the internal standard. Amino acid analysis was performed on Agilent 1260 HPLC (Agilent Technologies, Germany) equipped with an UV-Vis detector using automated ninhydrin derivatization.

### Solubility and aggregation analysis

Prior to analysis, a series of universal Britton-Robinson (ABP) buffers was prepared by mixing 20 mM acetic acid, 20 mM phosphoric acid and 20 mM boric acid and adjusting pH with 5 M NaOH. The ionic strength was adjusted with NaCl to conductivity that corresponds to the conductivity of 50 mM or 500 mM NaCl. Peptide libraries were thoroughly resuspended in autoclaved MilliQ water to the final concentration of 5 mg/ml. The peptide suspensions were diluted into the corresponding ABP buffer to the final concentration of 0.5 mg/ml and incubated for 30 min with mild shaking (500 rpm) at room temperature. After incubation, the insoluble fraction was separated by centrifugation at 21,300 *g* for 30 min at 4 ℃, and the supernatant (cleared sample) was subsequently used for solubility and aggregation analyses.

For all peptide libraries except 8E library, the amount of soluble peptides was estimated by measuring absorption of peptide bonds at 215 nm on a Nanodrop 2000 (Thermo Fisher Scientific, USA).

Due to its low solubility, the relative solubility of 8E peptide library was estimated with a fluorescamine assay(50) in a 96-well plate format. In brief, 90 μl of the peptide library solution was mixed with 30 μl of 3 mg/ml fluorescamine in DMSO, and the resulting mixture was incubated for 30 min in the dark at room temperature in a dark-bottom 96-well plate. The fluorescence was then recorded on Spark microplate reader (Tecan, Switzerland) using the following parameters: excitation at 365 nm, emission at 470 nm. 0.5 mg/ml solution of 8E peptide library in DMSO was used as a standard. All solubility measurements were conducted in technical duplicate for every library at every pH at two ionic strengths.

The relative amount of soluble aggregates was estimated by size-exclusion chromatography. A 100 μl aliquot of clarified sample (following centrifugation) was loaded onto a Superdex 75 Increase 10/300 GL column (Cytiva, USA) that was pre-equilibrated with two bed volumes of the corresponding 20 mM ABP buffer with either 50 or 500 mM NaCl. The peptides were eluted from the column isocratically on NGC Quest Plus System (Bio-Rad, USA), by one bed volume of the buffer at 0.5 ml/min flow rate at room temperature, and the eluted peptides were detected by absorption at 215 nm. The aggregation analysis was performed by quantifying the relative intensity underneath the peaks at ∼8 or ∼14 ml elution volume in the acquisition software (ChromLab Software v. 6.1). Size exclusion chromatography runs were conducted once for every library at every pH at two ionic strengths.

### Secondary structure analysis

The CD spectra were recorded using a Chirascan-plus spectrophotometer (Applied Photophysics, UK) over the wavelength range 190–260 nm in steps of 1 nm with an averaging time of 1 s per step. Cleared samples at 0.2 mg/ml nominal concentration in 1 mm path-length quartz cells were placed into a cell holder, and spectra were recorded at room temperature. The CD signal was obtained as ellipticity in units of millidegrees, and the resulting spectra were averaged from two scans and buffer-spectrum subtracted. All CD measurements were performed twice, and the resulting ratio of ellipticity at 222 nm to ellipticity at 200 nm was averaged and used to estimate the secondary structure content in peptide libraries.

To estimate the effect of pH on secondary structure, 5 mg/ml suspensions of peptide libraries in Milli-Q water were diluted into 10 mM ABP buffers at pH 3.0, 5.0, 7.4, 9.0, and 11.0 to the final nominal concentration of 0.2 mg/ml, gently mixed at room temperature for 30 min, and then centrifuged at 21,300 *g* for 15 min at 4 °C in order to remove insoluble fraction.

To estimate the effect of 2,2,2-trifluoroethanol on secondary structure, 5 mg/ml suspensions of peptide libraries in MilliQ water were diluted in a series of 10 mM ABP (pH 7.4) supplemented with 0–90% (v/v) 2,2,2-trifluoroethanol.

To estimate the effect of buffer on secondary structure, 5 mg/ml suspensions of peptide libraries in MilliQ water were diluted in 10 mM ABP (pH 7.4) or 10 mM Tris-HCl (pH 7.4).

### Infrared spectroscopy

The IR spectrum of 8E+Asp,Glu library was measured on Nicolet™ iS50 FTIR spectrometer (Thermo Fisher Scientific, Waltham, MA, USA), using a standard mid-IR source, KBr beamsplitter and a nitrogen-cooled MCT detector (2 cm^−1^ spectral resolution, Happ-Genzel apodization function, and 1000 scans) in the 4000–600 cm^−1^ range. The cell compartment was purged by dry nitrogen during the measurement. The sample was dissolved in a 50 mM PBS buffer (5 mg/ml nominal concentration). The CaF_2_ BioCell with a 8 μm path length (BioTools, Inc.) was used for the Fourrier-transform IR spectrum measurement. Spectral data treatment (GRAMS/AI) included solvent scan subtraction and background correction (using a linear function). Second derivative of the IR spectrum was calculated using the *Savitzky*-*Golay* numerical algorithm.

### Isolation and analysis of total, aggregate and monomer fractions from 8E+Asp,Glu library

The 800 μl aliquot of cleared 0.5 mg/ml 8E+Asp,Glu library solution in 10 mM ABP (pH 7.4) was loaded onto a Superdex 75 Increase 10/300 GL column (Cytiva, USA) that was pre-equilibrated with two bed volumes of 10 mM ABP buffer (pH 7.4). The peptides were eluted from the column by one bed volume of the buffer at 0.5 ml/min flow rate at room temperature, and the eluted peptides were detected by absorption of peptide bonds at 215 nm. The aggregate (∼8.5 ml) and monomer (∼13.5 ml) fractions were isolated and concentrated to 0.2 mg/ml using Amicon Ultra centrifugal filters (3 kDa MWCO). The total fraction was prepared by direct dilution of 0.5 mg/ml 8E+Asp,Glu library solution with 10 mM ABP (pH 7.4) to 0.2 mg/ml. The secondary structure analysis and HPLC amino acid analysis of total, aggregate and monomer fractions was performed as described above. The collection of the total, aggregate, and monomer fractions from the 8E+Asp,Glu library and the HPLC amino acid analysis on those fractions was conducted in technical triplicate. CD measurements and secondary structure analysis were conducted in technical duplicate.

The mass photometry data were collected using a TwoMP system (Refeyn, UK)(51). Prior to analysis, total, aggregate and monomer fractions were diluted with 10 mM Tris-HCl (pH 7.4), 50 mM NaCl to the final concentration of 20 μg/ml (∼8 μM). 17.5 µl of 10 mM Tris-HCl (pH 7.4), 50 mM NaCl was placed in a well of glass casket (Refeyn, UK) to focus the objective and subsequently mixed with 2.5 μl of the sample to achieve the final concentration of 2.5 μg/ml (∼1 μM). Mass photometry data was acquired and analyzed using AcquireMP and DiscoverMP software (Refeyn, UK), respectively. Mass calibration was performed using 10 nM bovine serum albumin (BSA) in 10 mM Tris-HCl (pH 7.4), 50 mM NaCl as a standard. The mass photometry analysis was performed once.

The morphology of monomer, aggregate and total fractions of 8E+Asp,Glu peptide library was studied by transmission electron microscopy (TEM) using negative staining. Prior to analysis, the 800 μl aliquot of cleared 0.5 mg/ml 8E+Asp,Glu library solution in 10 mM Tris (pH 7.4) was resolved into monomer and aggregate fractions according to the aforementioned protocol. The total fraction was prepared by direct dilution of 0.5 mg/ml 8E+Asp,Glu library solution with 10 mM Tris (pH 7.4) to 0.2 mg/ml. Additionally, the total fraction of 8E+Lys,Arg,Asp,Glu peptide library was used as a reference library with no observable aggregation. For the analysis, 0.2 mg/ml samples in 10 mM Tris (pH 7.4) were diluted with buffer to 20 μg/ml final concentration. The parlodion-carbon coated, 200 mesh copper grids (SPI supplies, USA) were glow discharged with GLOQUBE plus instrument (Quorum Technologies, UK) and soaked with 20 µl of the sample solution for 1 min at room temperature. The excess sample was soaked away with filter paper (Whatman), and the grids were subsequently stained two times with 20 µl of 2% (w/v) freshly filtered uranyl acetate in water for 1 min at room temperature. The grids were dried overnight at room temperature and examined in JEM-2100PLUS transmission electron microscope (JEOL, Japan) equipped with TemCam-XF416 CMOS camera (TVIPS, Norway) that was operated at 200 kV.

The stability of 8E+Asp,Glu aggregate fraction was assessed by CD spectroscopy measured upon titration with guanidine hydrochloride and temperature-dependent CD spectroscopy. The isolated aggregate fraction (prepared in the same way as described above) in 10 mM ABP (pH 7.4) was concentrated to 0,1 mg/ml using Amicon Ultra centrifugal filters (3 kDa MWCO). The ECD spectra were measured using the Jasco 1500 spectropolarimeter equipped with the Peltier holder PTC-517. Temperature dependence of CD spectra was measured in spectral range 200-280 nm from 10 °C to 90 °C with the step of 10 °C in 0.5 mm path length quartz cell (scanning speed of 10 nm/min, response time of 8 seconds, scanning step 0.5 nm, 1 accumulation). For the guanidine hydrochloride titration experiment, a concentration range of 1–2 M was employed. After baseline correction, the final spectra were expressed as molar ellipticity θ (deg·cm^2^·dmol^-1^) per residue.

To support the CD results and prove the stability of 8E+Asp,Glu assemblies, isolated fraction (prepared in the same way as described above) in 10 mM ABP (pH 7.4) was split in 4 aliquots (900 ml) and incubated separately at 20, 70, 80 and 90 °C for 1 hour prior to analytical SEC. 800 μl aliquot of sample was loaded onto a Superdex 75 Increase 10/300 GL column (Cytiva, USA). The peptides were eluted from the column isocratically on NGC Quest Plus System (Bio-Rad, USA), by one bed volume of the buffer (10mM ABP, pH 7.4) at 0.5 ml/min flow rate at room temperature, and the eluted peptides were detected by absorption at 215 nm.

### Metal chelation assay using Fura-2 fluorescent reporter

Metal chelation assays were performed by monitoring the fluorescence of the metal-sensitive dye Fura-2 in the presence of various divalent metal ions and competing peptide ligands. The assay was conducted in a 384-well microplate format, with all measurements performed in triplicate.

Each well contained a total volume of 21.08 µl. The final well composition consisted of 10 µl of metal ion solution in 100 mM HEPES buffer at pH 7.4, 10 µl of Na₅.Fura-2 solution in the same buffer (both titrated to pH 7.4 using NaOH), and 1.08 µl of a peptide library stock solution in 10 mM DMF (final concentration of DMF was 5.3% v/v).

The metal ion concentration (Fe^2+^, Ca^2+^, Zn^2+^, Ni^2+^, and Co^2+^) in each well was 16.6 µM, while the Fura-2 concentration was 2.2 µM. Peptides were added at a final concentration of 510 µM to assess their ability to competitively bind metal ions and displace Fura-2 from the complex. Notably, the Fe^2+^ stock appeared partially oxidized, as indicated by an EDTA titration, which affected the assay only by lowering the effective Fe^2+^ concentration, since Fe^3+^ is not detectable by Fura-2.

Four different peptides libraries were tested for their metal chelation ability: 10E (assumed MW 2474.24), 10KR (MW 2574.56), 12DEKR (MW 2656.94), and 19F (MW 3008.93). All peptides were added to achieve the same final molar concentration of 510 µM, with corresponding stock concentrations prepared in the range of 1.3 to 1.5 mg/ml depending on peptide molecular weight.

Fluorescence changes of the Fura-2–metal complexes at E_x_ 363 nm, E_m_ 512 nm for Fe^2+^ and Co^2+^, and E_x_ 335 nm, E_m_ 505 nm for Ca^2+^, Zn^2+^ and Ni^2+^, were monitored following peptide addition to determine competitive chelation behavior. Fractional metal ion chelation (Chelation %) was calculated by linear normalization as:

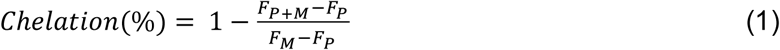

Where *F_P_*_+*M*_ is the fluorescence intensity of the mixture containing Fura-2, peptide library, and metal ions; *F_P_* is the intensity of Fura-2 with the peptide library alone (100 % chelation control); and *F_M_* is the intensity of Fura-2 with the metal ions alone (0 % chelation control). Plotted values show medians of triplicate measurements, with standard deviations as error bars.

### Radius of gyration on predicted peptide structures

Fifty thousand (50,000) unique and random 25-mer peptide sequences were generated *in-silico* by uniformly sampling the amino acid alphabets of each library (8E, 8E+Asp,Glu, 8E+Lys,Arg, 8E+Lys,Arg,Asp,Glu and the full canonical alphabet of 20 amino acids). The heavy atom coordinates of all two hundred fifty thousand (250,000) peptide sequences were predicted using ESMFold-v1(32). These structures were used to calculate the radius of gyration (GROMACS 2024.2). All histograms and kernel density estimations were calculated using Python v3.12.0 and SciPy v1.13.0.

## Data availability

To facilitate resynthesis, all product codes and loadings for Fmoc-protected amino acids used for solid phase peptide synthesis are reported as Supplemental Data S1. To facilitate reanalysis, raw CD spectral data, solubility data, and amino acid analysis are supplied as Supplemental Data S2. Computational results including predicted radius of gyration and secondary structure for 50,000 random sequences for 5 alphabets are supplied in Supplemental Data S3. The scripts used for ESMfold predictions and structural analyses are available here: https://github.com/Singharoy-Group/ESMFold_Application.git

## Acknowledgements

We thank Dr. Radko Souček and Dr. Kateřina Nováková for their technical support during the quality control of the peptide libraries. We thank John Cava for assistance with installation of ESMFold on compute clusters. We thank Markéta Pazderková for help with the interpretation of the infrared spectrum. In addition, we would like to thank Prof. Stephen Freeland for helpful discussions about this work. This work was supported by a Research Grant from HFSP https://doi.org/10.52044/HFSP.RGEC272023.pc.gr.168579. S.D.F. acknowledges support from the NIH Director’s New Innovator Award (DP2-GM140926), a Cottrell Scholar Award, and a Sloan Fellowship for support. E.M.S. thanks the Program in Molecular Biophysics training grant (NIH-T32GM135131). M.M. and K.H. acknowledge CMS CF Biophysical techniques of CIISB, Instruct-CZ Centre, supported by MEYS CR Infrastructure project LM2018127 and European Regional Development Fund-Project, “UP CIISB” (No. CZ.02.1.01/0.0/0.0/18_046/0015974). Computational resources were provided by the e-INFRA CZ project (ID:90254), supported by the Ministry of Education, Youth and Sports of the Czech Republic.

