## Supporting Information for "Prebiotically Plausible Peptides can Self-assemble into β-rich Assemblies"

**This PDF file includes:**

Supplemental Figures S1 to S15
Supplemental Table S1

**Figures**

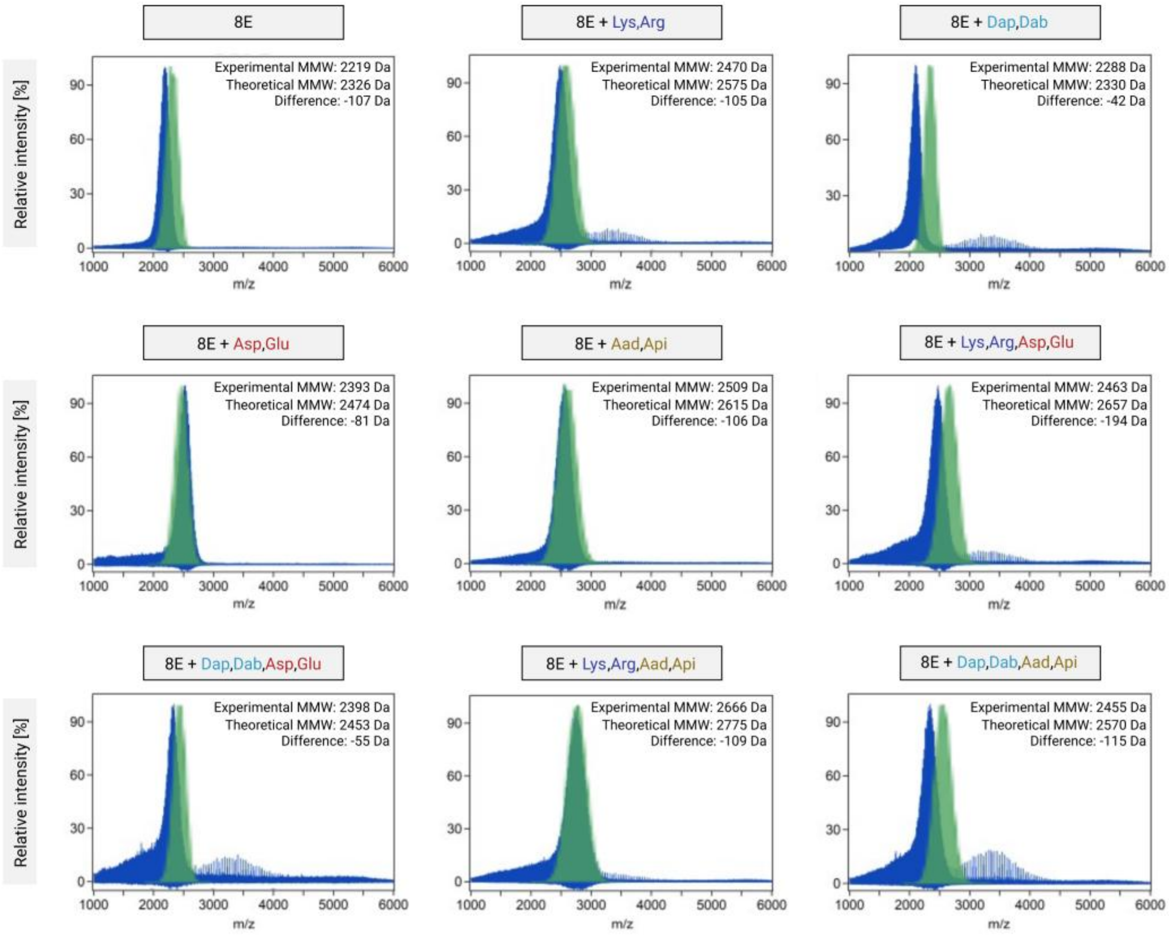


**Supplemental Figure S1. Molecular weight distribution of nine indicated peptide libraries as assessed by MALDI.** MALDI spectra (blue) were obtained on 10 mg/ml solutions of peptide libraries dissolved in acetonitrile:water (1:1) mixture. The theoretical distributions (green) were obtained from 10,000,000 randomly generated sequences with equimolar amino acid distribution. For each library, the mean molecular weight (MMW) observed experimentally and the theoretical mean molecular weight are reported.

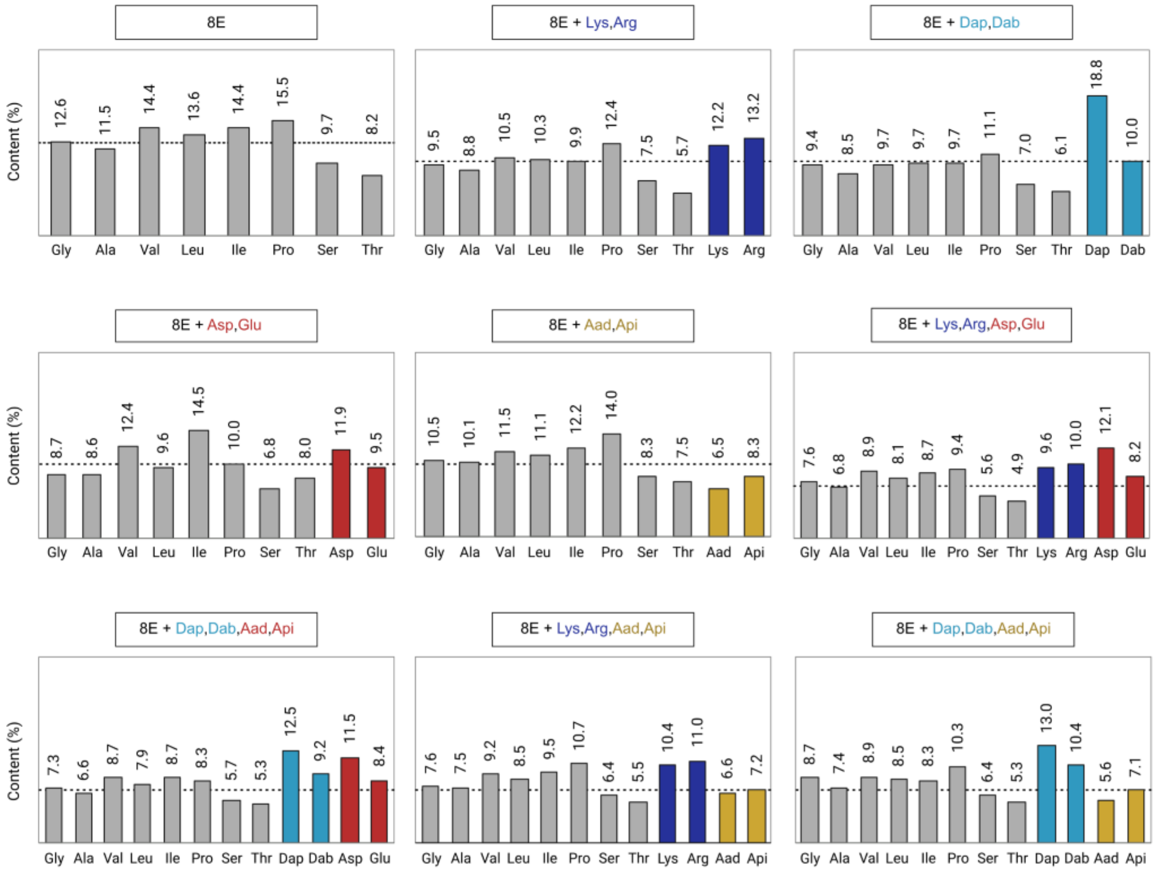


**Supplemental Figure S2. The amino acid composition of nine indicated peptide libraries as assessed by HPLC amino acid analysis.** Theoretical amino acid distributions (dashed line) represent equimolar distribution of amino acids for each library and experimental distributions (bars) reflect integrated ion intensities from HPLC chromatograms.

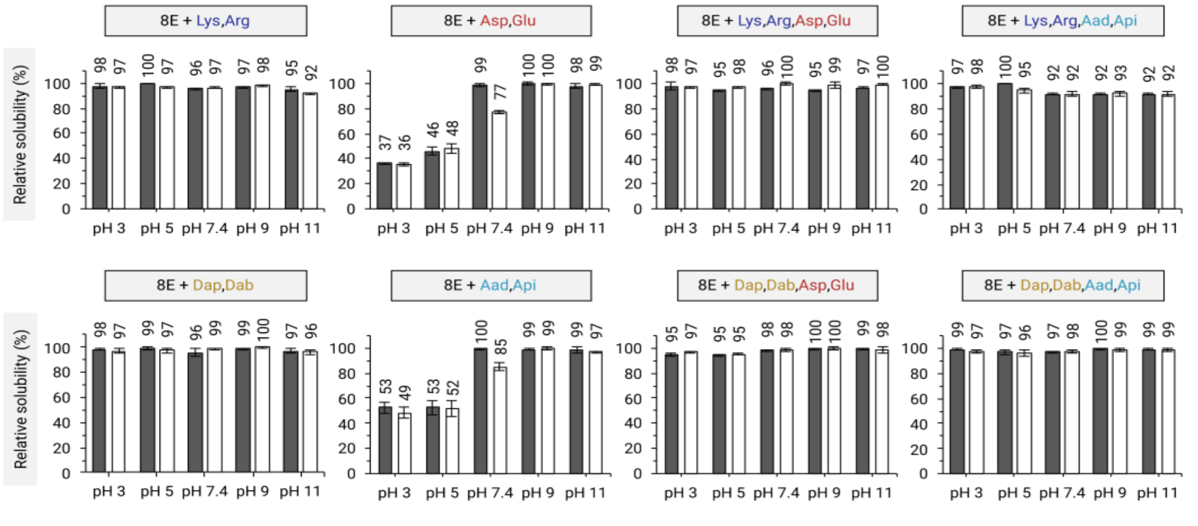


**Supplemental Figure S3. The relative solubility of eight peptide libraries.** Solubility of the indicated libraries was measured in 20 mM ABP buffer at different pH's and ionic strengths by absorption of peptide bonds at 215 nm. Grey bars correspond to medium (50 mM NaCl) and white bars correspond to high (500 mM NaCl) ionic strength. Error bars represent standard deviations from duplicates with the exception of 8E+Asp,Glu library (triplicates). The data for 8E+Asp,Glu were taken from Ref. (20) in the main text.

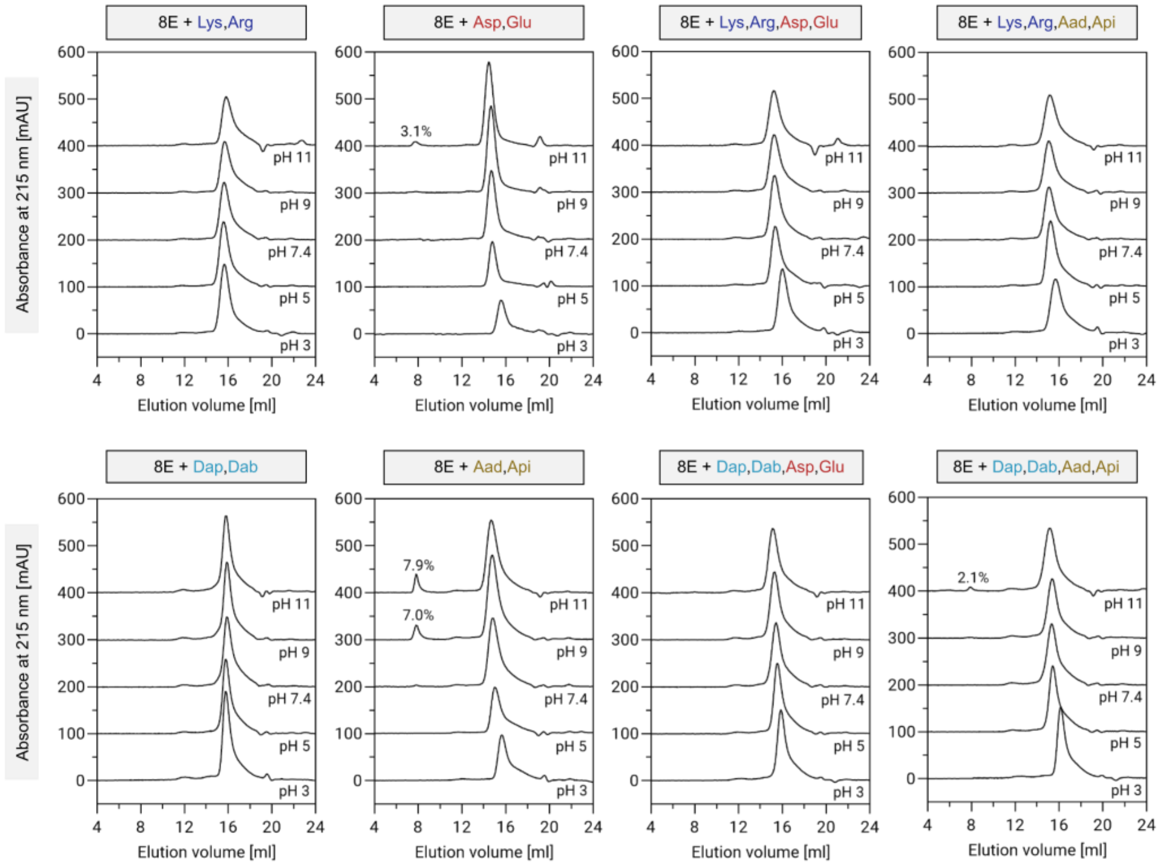


**Supplemental Figure S4. The aggregation propensity of eight peptide libraries at high ionic strength.** Aggregation propensity of the indicated libraries was measured in 20 mM ABP buffer at different pH's and at high ionic strength (500 mM NaCl) with size-exclusion chromatography (SEC). SEC chromatograms report absorbance at 215 nm as a function of elution volume. The data for 8E+Asp,Glu library were taken from Ref. (20) in the main text. The respective chromatograms were shifted along the y axis by 100 mAU for easy comparison.

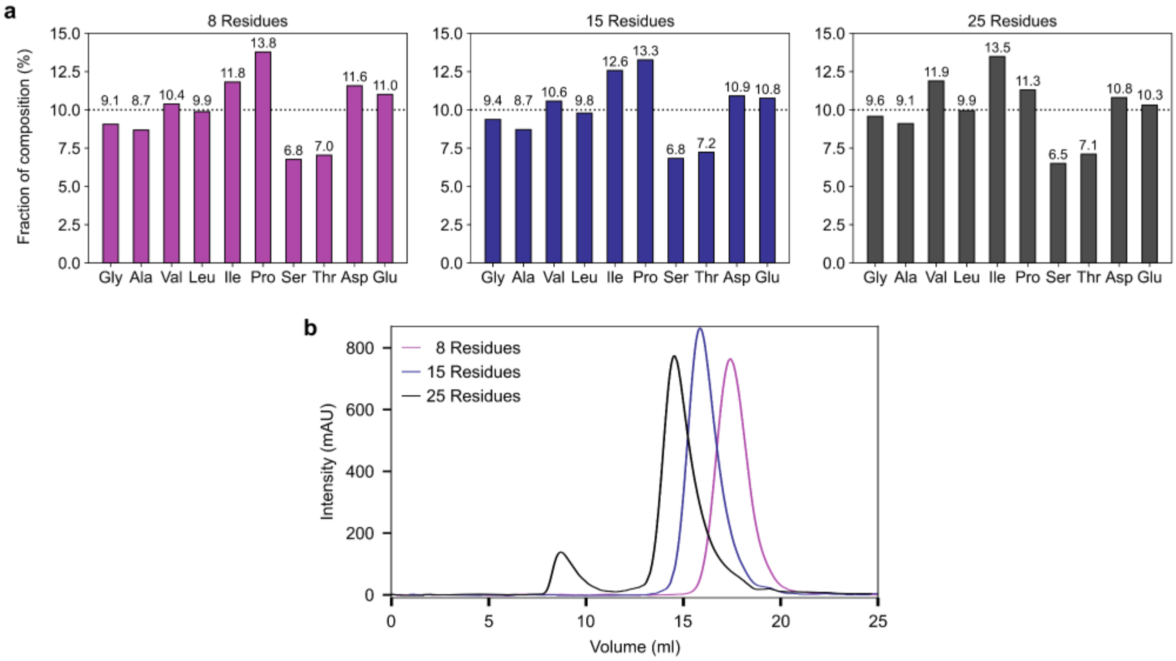


**Supplemental Figure S5. Length dependence of the aggregation propensity of 8E+Asp,Glu library.** (**a**) Amino acid composition of additionally synthesized 8, 15, and 25-mer 8E+Asp,Glu libraries as assessed by HPLC amino acid analysis. Theoretical amino acid distributions (dashed line) represent equimolar distribution of amino acids for each library, and experimental distributions (bars) reflect integrated ion intensities from HPLC chromatograms. (**b**) Aggregation propensity of the indicated 8E+Asp,Glu libraries was measured in 20 mM ABP buffer (pH 9.0) at medium ionic strength (50 mM NaCl) with size-exclusion chromatography (SEC). SEC chromatograms report absorbance at 215 nm as a function of elution volume.

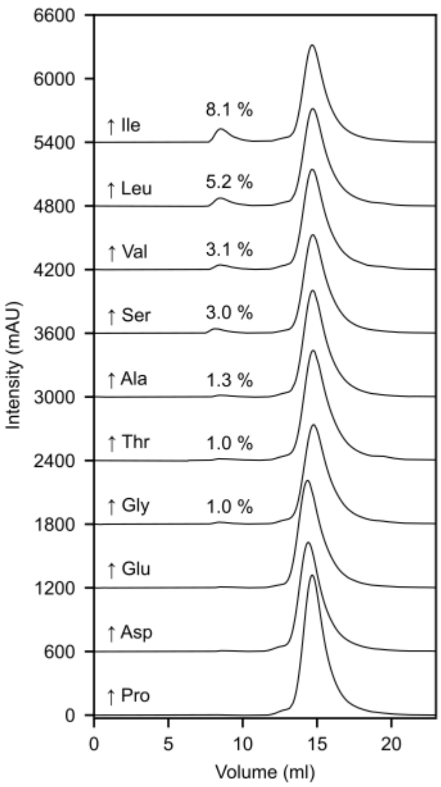


**Supplemental Figure S6. The aggregation propensity of 8E+Asp,Glu libraries with individually skewed amino acid ratios.** Aggregation propensity of the indicated 8E+Asp,Glu libraries was measured in 20 mM ABP buffer (pH 9.0) at medium ionic strength (50 mM NaCl) with size-exclusion chromatography (SEC). SEC chromatograms report absorbance at 215 nm as a function of elution volume. See Supplemental Table 1 for the ratios of the amino acids in the mixtures. The respective chromatograms were shifted along the y axis by 600 mAU to facilitate comparison.

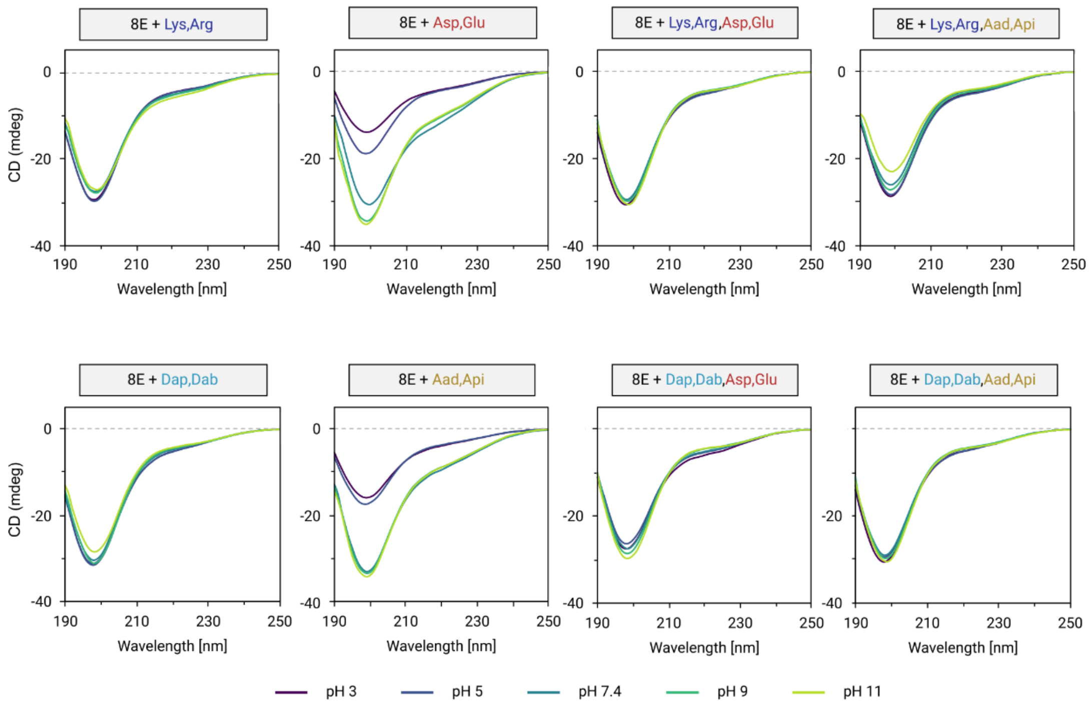


**Supplemental Figure S7. The secondary structure of eight peptide libraries.** Secondary structure of the indicated libraries was measured in 10 mM ABP buffer at different pH's with far-UV circular dichroism (CD) spectroscopy. Far-UV CD spectra are representative from duplicates. The data for 8E+Asp,Glu library were taken from Ref. (20) in the main text.

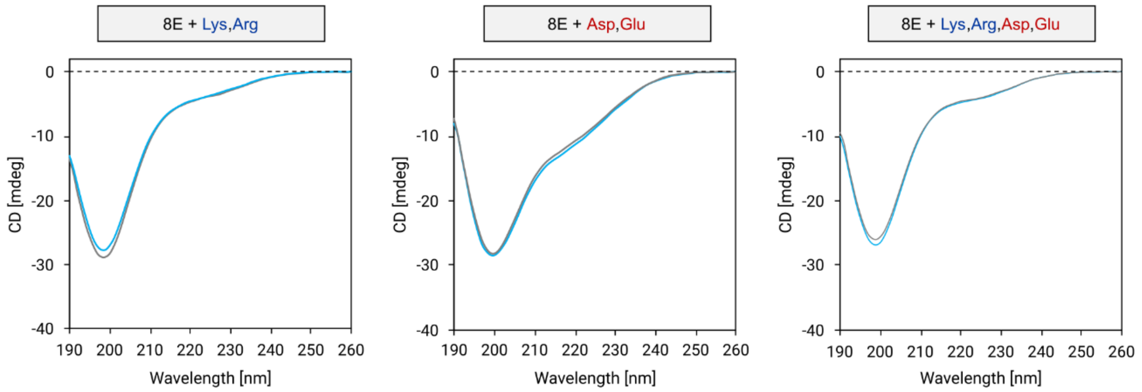


**Supplemental Figure S8. The effect of buffer on peptide library secondary structure.** The secondary structure of three indicated peptide libraries in 10 mM Tris-HCl (pH 7.4, grey) and in 10 mM ABP (pH 7.4, blue) as assessed by far-UV circular dichroism (CD) spectroscopy. No significant differences are observed between the two buffers.

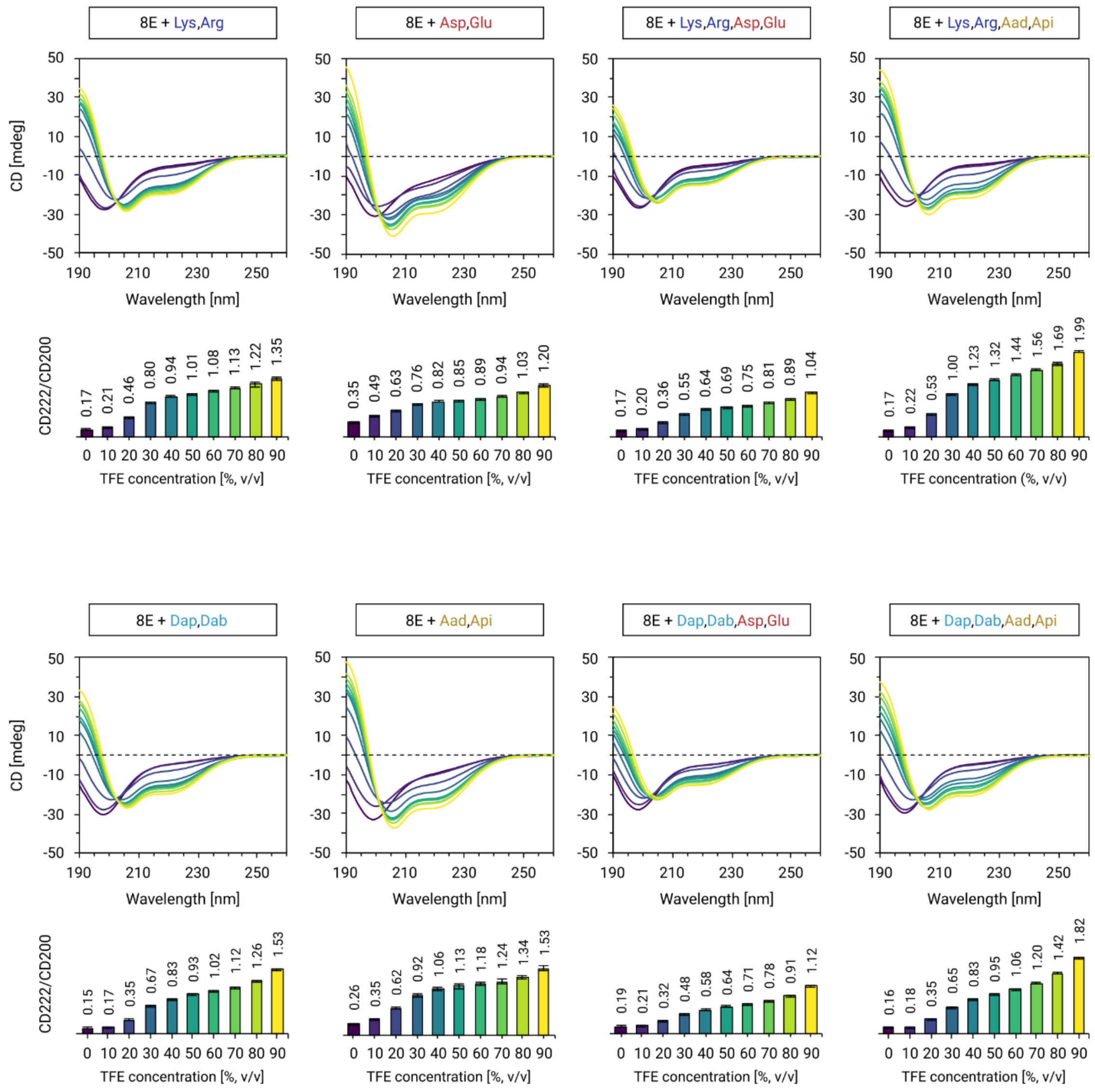


**Supplemental Figure S9. Induction of secondary structure with 2,2,2-trifluoroethanol (TFE).** The secondary structure of eight peptide libraries at different TFE concentrations as assessed by far-UV circular dichroism (CD) spectroscopy. Far-UV CD spectra were measured in 10 mM ABP (pH 7.4) with 0–90% (v/v) TFE and are representative from duplicates. The approximate secondary structure content is estimated as the ratio of the ellipticities at 222 nm and 200 nm as shown in the bar charts. Error bars represent standard deviations from duplicates. The data for 8E+Asp,Glu library were taken from Ref. (20) in the main text.

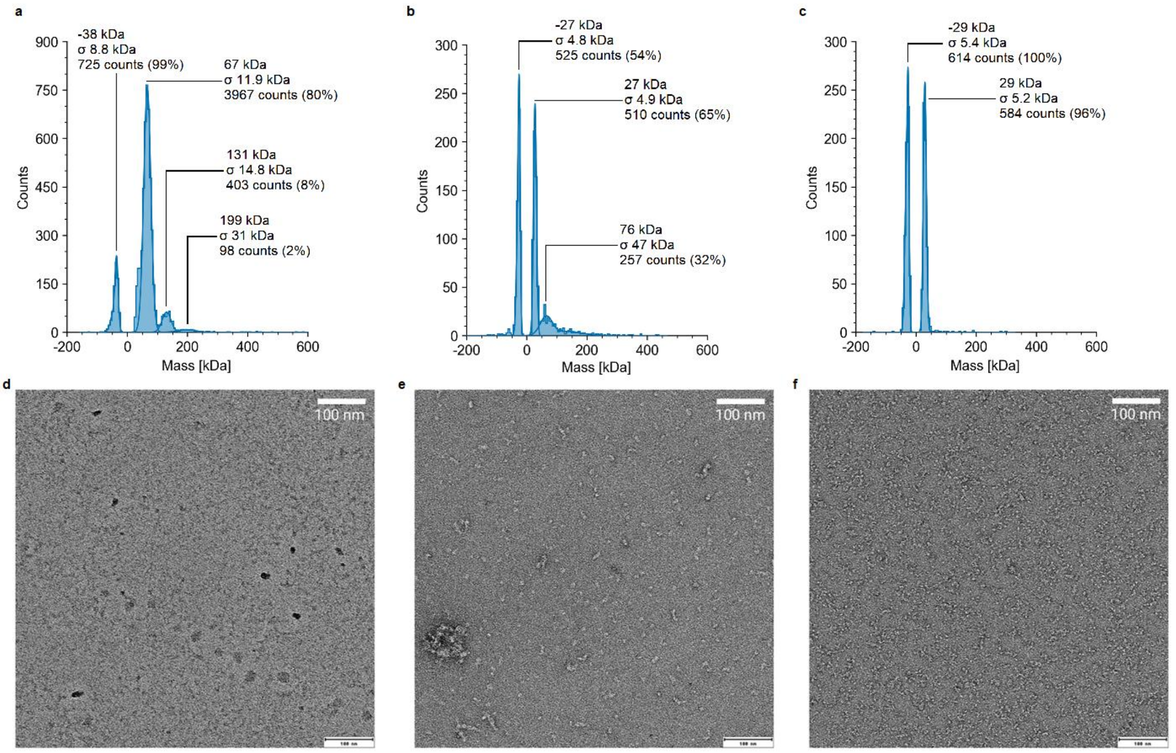


**Supplemental Figure S10. Analysis of 8E+Asp,Glu with mass photometry and transmission electron microscopy.** The molecular weight distribution of 8E+Asp,Glu library (and its SEC-resolved fractions) in 10 mM Tris-HCl (pH 7.4), 50 mM NaCl as assessed by mass photometry. Mass histograms of: (**a**) BSA standard; (**b**) total fraction and (**c**) monomer fraction. Transmission electron micrographs of (**d**) total fraction of 8E+Asp,Glu,Lys,Arg library; (**e**) total fraction; (**f**) monomer fraction 8E+Asp,Glu library. The electron micrographs were obtained on samples with 20 µg/ml concentration in 10 mM Tris-HCl (pH 7.4).

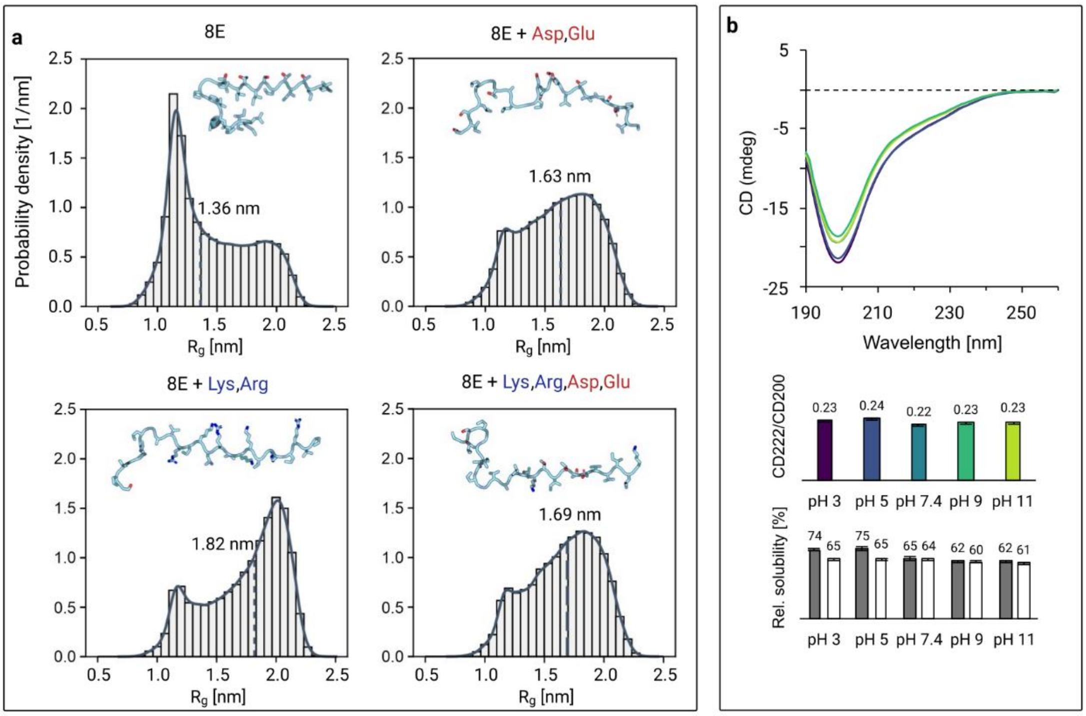


**Supplemental Figure S11. A large language model prediction of library structure compaction.** (**a,c**) The structures of 50,000 unique and randomly generated 25-mer peptide sequences were predicted using ESMFold (ref. 32 in the main text) for each 8E, 8E+Asp,Glu, 8E+Lys,Arg and 8E+Lys,Arg,Asp,Glu libraries. (**a**) Distribution of the radius of gyration calculated with GROMACS 2024.2. Representative peptide structures centered at the distributions' medians (dashed lines) are shown above. (**b**) Secondary structure and solubility of 8E library at various pH's.

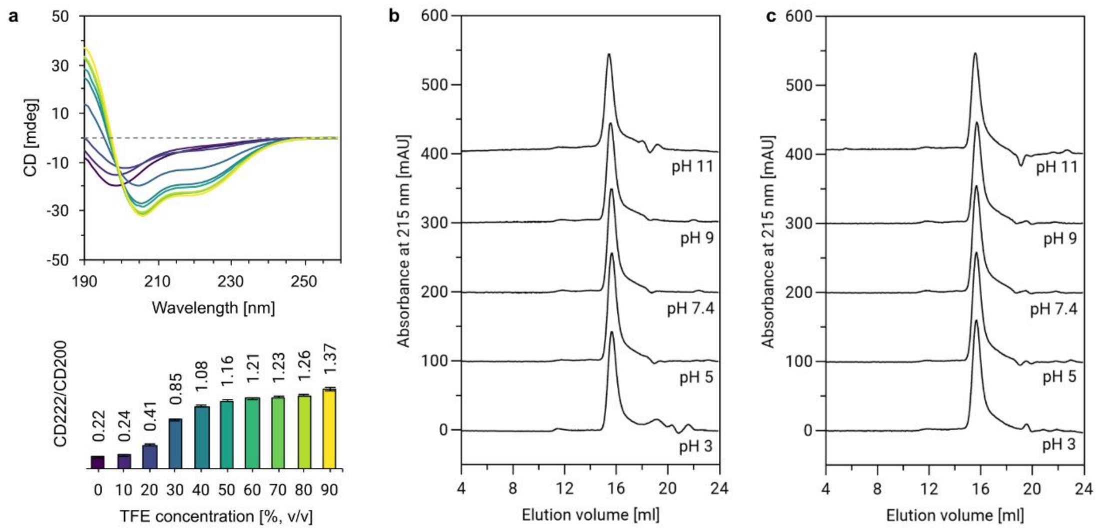


**Supplemental Figure S12. Additional assays on the hydrophobic 8E peptide library.** (**a**) The secondary structure of the 8E peptide library at different 2,2,2-trifluoroethanol (TFE) concentrations as assessed by far-UV circular dichroism (CD) spectroscopy. Far-UV CD spectra were measured in 10 mM ABP (pH 7.4) with 0–90% (v/v) TFE and are representative from duplicates. The secondary structure content is estimated as the ratio of the ellipticities at 222 nm and 200 nm as shown in the bar charts. Error bars represent standard deviations from duplicates. (**b,c**) Aggregation propensity of the 8E peptide library was measured in 20 mM ABP buffer at different pH's and at medium ionic strength (50 mM NaCl, **b**) and at high ionic strength (500 mM NaCl, **c**) with size-exclusion chromatography (SEC). SEC chromatograms report absorbance at 215 nm as a function of elution volume.

**Supplemental Tables**

**Supplemental Table S1. Amino acid composition of 8E+Asp,Glu libraries with individually skewed amino acid ratios.** Measured molar percent are displayed.

| | Libraries (enriched amino acid in bold) | | | | | | | | | |
|---|---|---|---|---|---|---|---|---|---|---|
| Library | 1. | 2. | 3. | 4. | 5. | 6. | 7. | 8. | 9. | 10. |
| Gly (%) | **15.71** | 10.89 | 10.59 | 10.01 | 10.98 | 9.74 | 10.09 | 9.99 | 9.86 | 10.10 |
| Ala (%) | 9.36 | **16.77** | 10.46 | 9.33 | 10.08 | 9.62 | 9.48 | 9.35 | 9.20 | 9.48 |
| Val (%) | 9.60 | 11.01 | **13.17** | 8.98 | 9.22 | 9.87 | 8.70 | 8.61 | 8.10 | 9.07 |
| Leu (%) | 10.69 | 10.52 | 10.64 | **15.47** | 10.69 | 10.62 | 10.10 | 10.21 | 9.90 | 10.21 |
| Ile (%) | 8.26 | 7.62 | 7.57 | 7.94 | **12.71** | 8.73 | 7.93 | 8.03 | 7.86 | 7.95 |
| Pro (%) | 11.98 | 11.19 | 13.37 | 12.05 | 11.72 | **16.51** | 13.09 | 13.27 | 14.38 | 11.88 |
| Ser (%) | 8.75 | 7.88 | 8.93 | 9.10 | 8.60 | 9.19 | **13.80** | 9.10 | 9.01 | 8.68 |
| Thr (%) | 7.63 | 7.15 | 7.66 | 8.32 | 7.49 | 7.60 | 7.99 | **12.82** | 8.47 | 8.07 |
| Asp (%) | 8.75 | 8.58 | 8.59 | 8.93 | 8.48 | 8.76 | 8.71 | 8.54 | **12.95** | 8.79 |
| Glu (%) | 9.28 | 8.39 | 9.02 | 9.87 | 10.02 | 9.37 | 10.13 | 10.06 | 10.26 | **15.75** |
